## Supplementary material for "Over-expression of the brassinosteroid gene *TaDWF4* increases wheat productivity under low and sufficient nitrogen through enhanced carbon assimilation": suppl figures

Suppl. Table 1: Percent Identity Matrix of TaDWF4 genes in wheat compared to the rice DWF4 (Os030227700)

|  |  | Percent identity | | | | | | | |
| --- | --- | --- | --- | --- | --- | --- | --- | --- | --- |
| Gene Locus | | TraesCS3D01G526400 | Os03g0227700-OsDWF4 | TraesCS4B01G234100 | TraesCS4A01G078000 | TraesCS4D01G235200 | TraesCS3A01G519000 | TraesCS3B01G586500 | TraesCS3D01G526300 |
| TraesCS3D01G526400 | | 100 | 80.52 | 83.96 | 84.43 | 84.2 | 89.86 | 87.97 | 90.07 |
| Os03g0227700-OsDWF4 | | 80.52 | 100 | 89.62 | 89.22 | 89.42 | 84.44 | 84.85 | 86.03 |
| TraesCS4B01G234100 | | 83.96 | 89.62 | 100 | 98.62 | 98.81 | 89.78 | 90.98 | 91.35 |
| TraesCS4A01G078000 | | 84.43 | 89.22 | 98.62 | 100 | 99.01 | 89.98 | 90.98 | 91.35 |
| TraesCS4D01G235200 | | 84.2 | 89.42 | 98.81 | 99.01 | 100 | 89.98 | 91.18 | 91.55 |
| TraesCS3A01G519000 | | 89.86 | 84.44 | 89.78 | 89.98 | 89.98 | 100 | 92.99 | 95.37 |
| TraesCS3B01G586500 | | 87.97 | 84.85 | 90.98 | 90.98 | 91.18 | 92.99 | 100 | 95.77 |
| TraesCS3D01G526300 | | 90.07 | 86.03 | 91.35 | 91.35 | 91.55 | 95.37 | 95.77 | 100 |

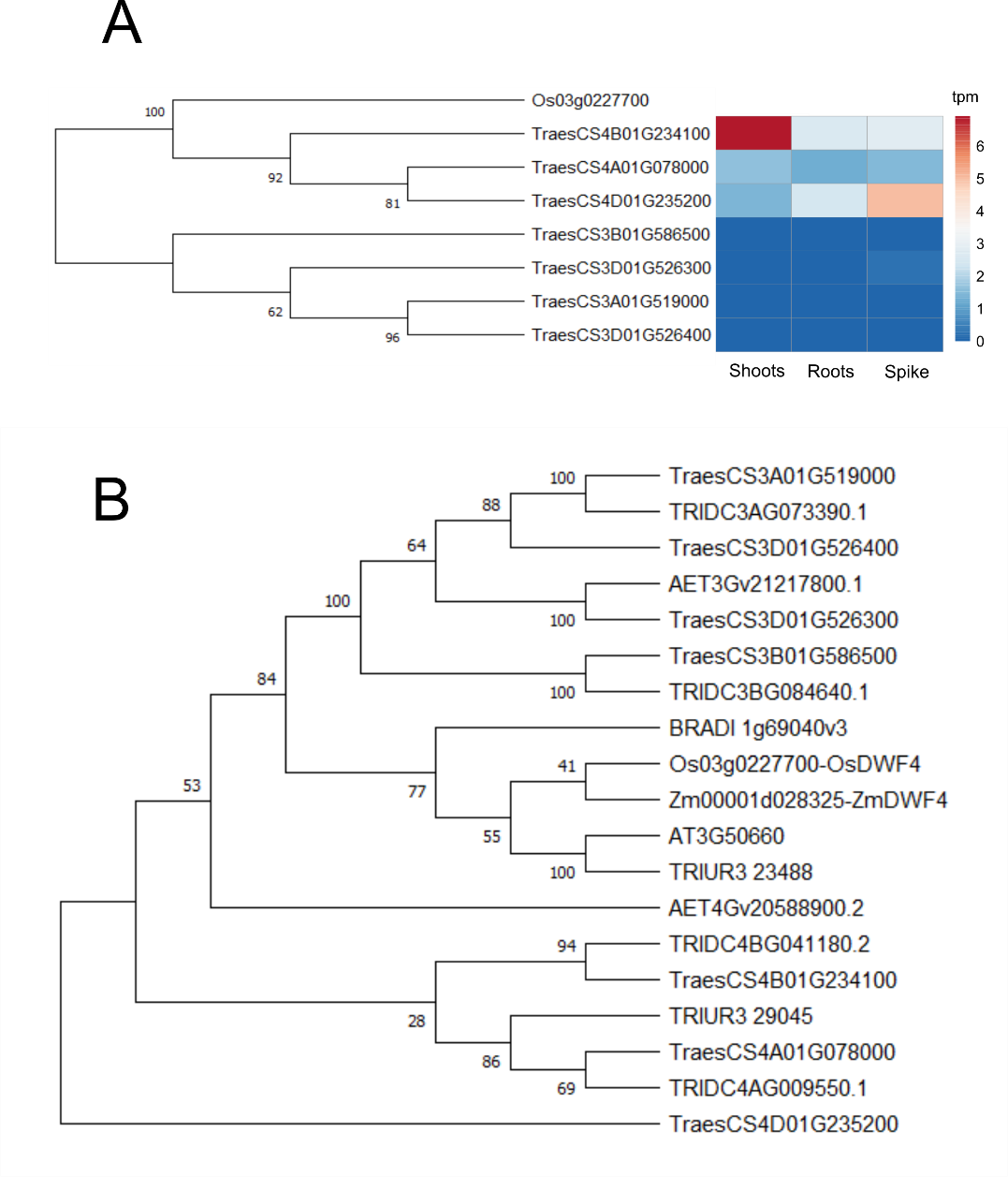

Suppl. Figure 1: Phylogenic tree DWF4 genes in wheat and it progenitors. A) Phylogenic tree and expression levels in three tissues of the seven putative DWF4 genes in wheat relative to the known DWF4 gene from rice. Values on the tree represent the boot strap value from 500 iterations of aligning the sequences. Color code of the expression correspond to corresponding gene models based on the RefSeq v1 wheat genome. Expression values shown via heatmap of the seven putative DWF4 genes in wheat cv. Chinese spring shown is tpm (transcripts per million). B) Phylogenitc tree of DWF4 genes from wheat, rice, maize, *T.uratu, A.tauschii* and *T. dicoccoides*.

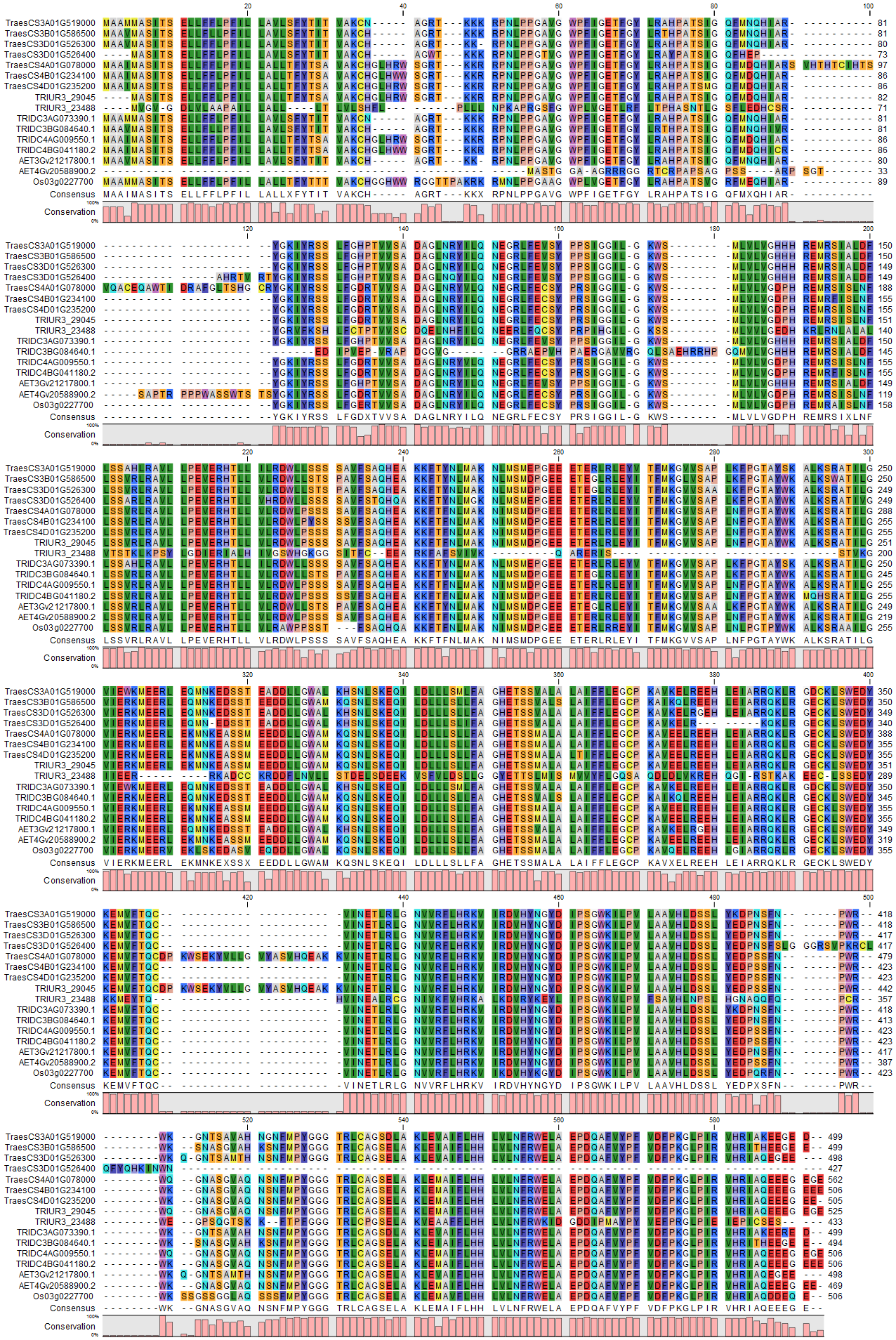

Suppl. Figure 2: Alignment of the amino acid sequences of the first isoform of each DWF4 homeologue from wheat and its progenitor species. Labels beginning with Treas are from bread wheat (refseq v1), TRUR are from *T.uratu,* AET from *A.tauschii*, TRIDC from *T. dicoccoides,* Os Oryza sativa.

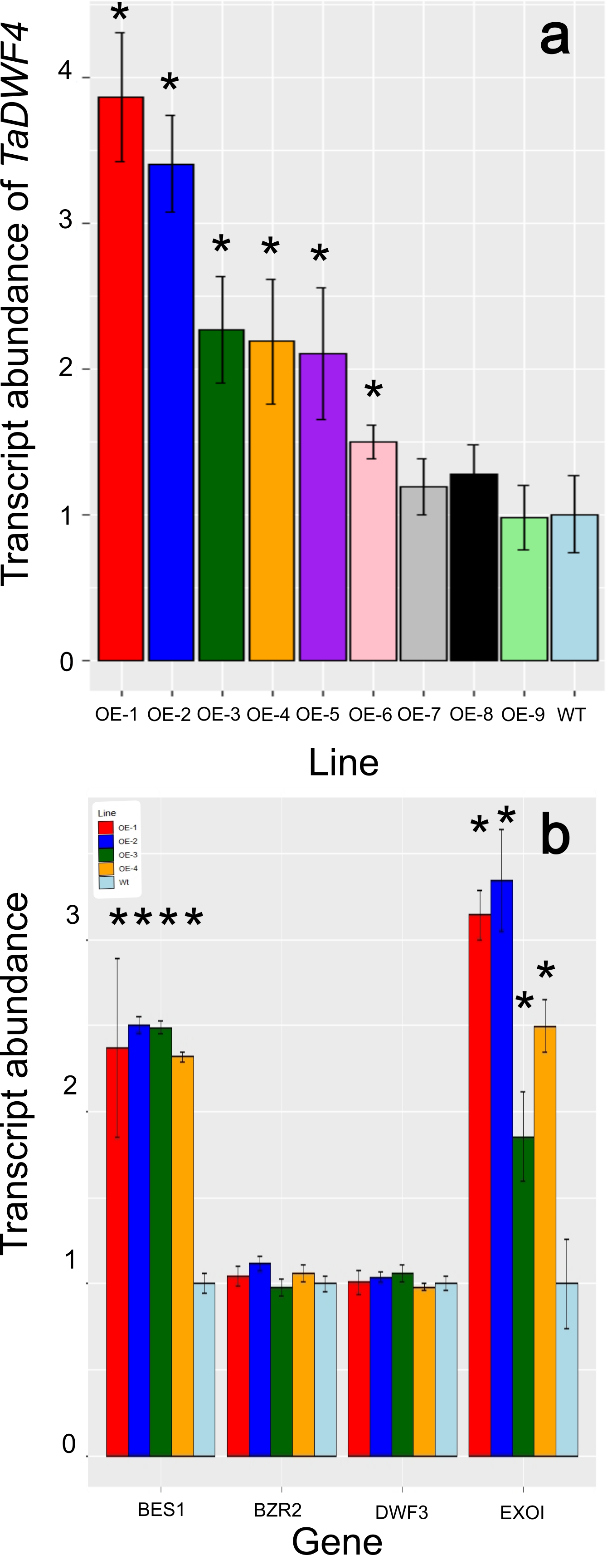

Suppl. Figure 3: Relative expression of *TaDWF4* and BR related genes as markers of BR levels in wheat over expression lines. A) *TaDWF4* expression in over expressing lines with a single copy T-DNA insertion. Expression shown is relative to *TaUbi*. B) Relative expression of *TaBES1, TaBZR1, TaDWF3 and TaEXOI* in the highest four *TaDWF4* over expressing lines with a single copy T-DNA insertion. Expression with SE is shown is relative to *TaUbi* from shoots of 14 day old plants. Significant differences are noted as * relative to WT expression (p val <0.05).

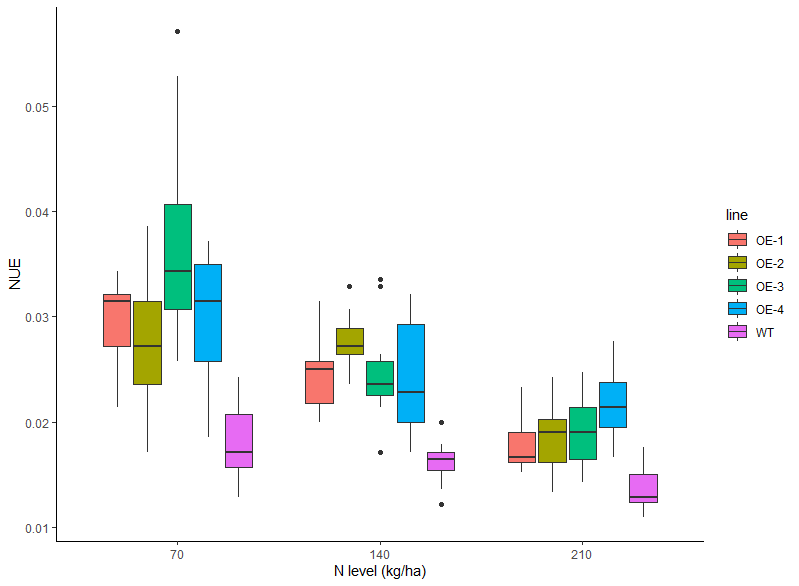

Suppl. Figure 4: NUE of *CYP90B-B* overexpressioin lines grown under three different N concentrations

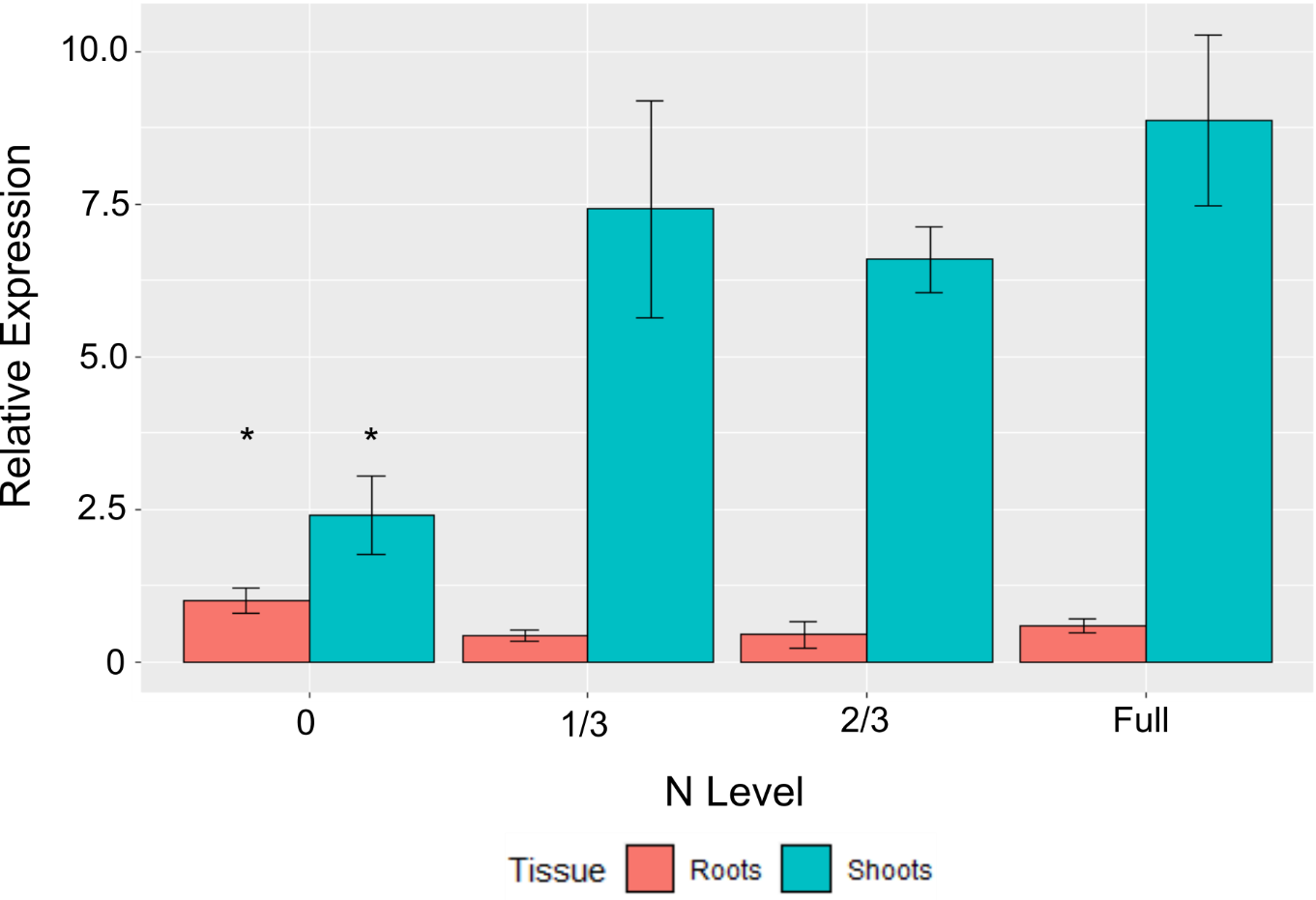

Suppl. Figure 5: Expression of *TaCYP90B* in Fielder wheat plants grown in hydroponics with various levels of N in solution roughly equivalent to 70, 140 and 210 kg/ha which equates to 1/3, 2/3 and Full N in hydroponics. Proportion of N is relative to the full strength solution. TaDWF4 expression is shown relative to the DWF4 expression grown with no N in the roots. Significant differences are noted as * (p val <0.05).

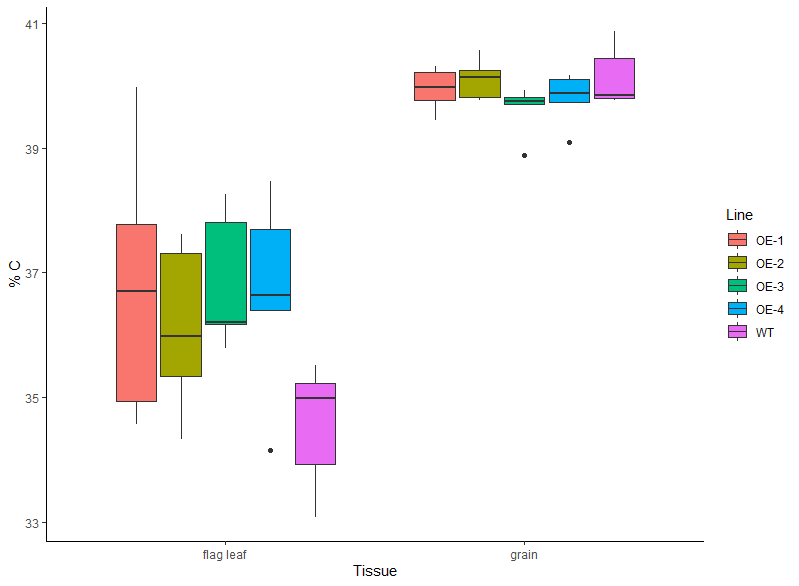

Suppl. Figure 6: Carbon content of flag leaves and grain.

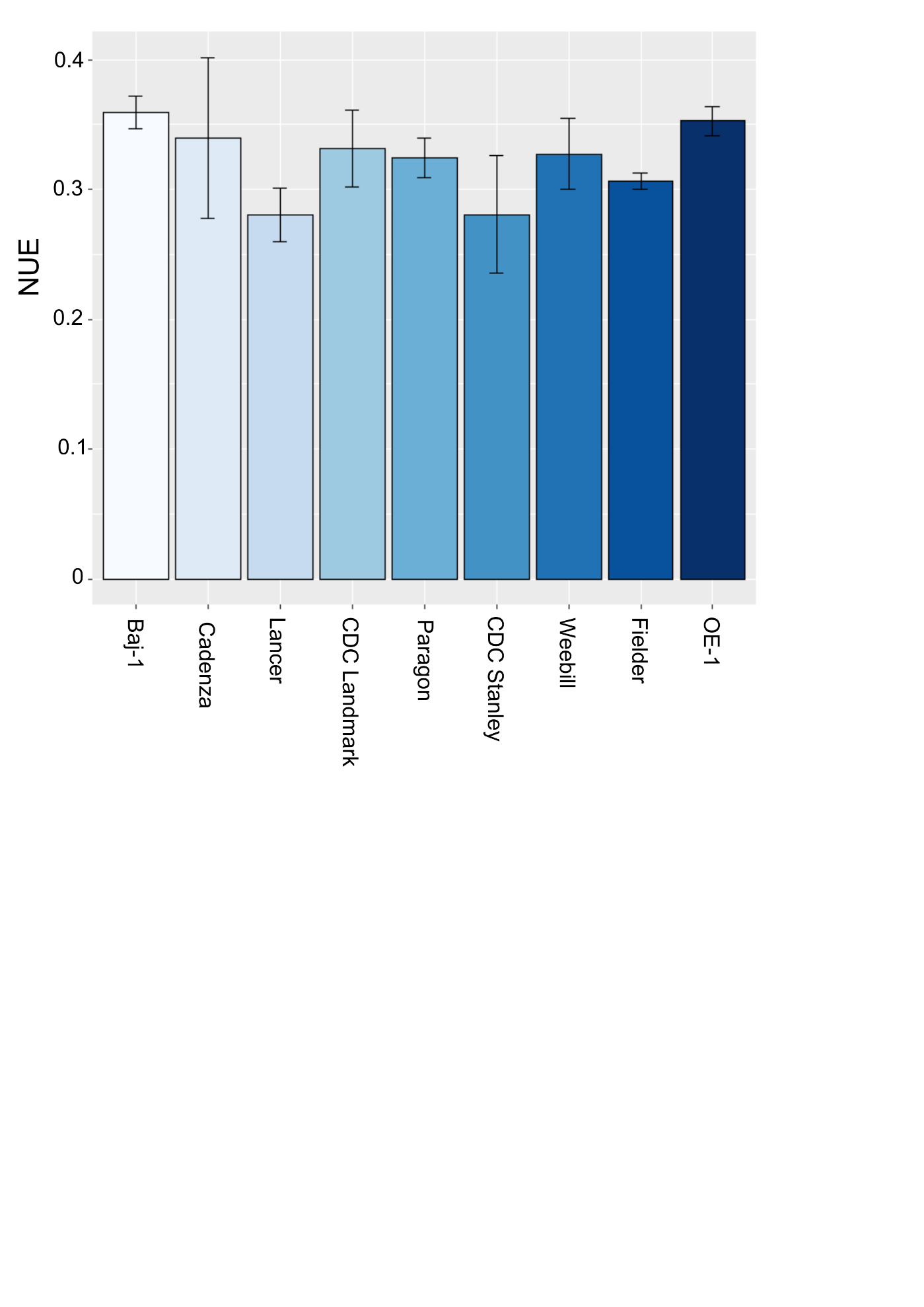

Suppl. Figure 7: NUE of spring wheat varieties grown on a low fertility soil supplemented with either 70 kg/ha (low) or 210 kg/ha (high). Data shown is the means of six plants for each treatment for the yield on low N divided by high N.
